# Maternal hypothyroidism in mice influences glucose metabolism in adult offspring

**DOI:** 10.1101/785030

**Authors:** Yasmine Kemkem, Daniela Nasteska, Anne De Bray, Paula Bargi-Souza, Rodrigo A. Peliciari-Garcia, Anne Guillou, Patrice Mollard, David J. Hodson, Marie Schaeffer

## Abstract

During pregnancy, maternal metabolic diseases and hormonal imbalance may alter fetal beta cell development and/or proliferation, thus leading to an increased risk for developing type 2 diabetes in adulthood. Although thyroid hormones play an important role in fetal endocrine pancreas development, the impact of maternal hypothyroidism on glucose homeostasis in adult offsprings remains poorly understood. Here, we show that when fed normal chow, adult mice born to hypothyroid mothers were more glucose-tolerant due to beta cell hyperproliferation and increased insulin sensitivity. However, following high fat feeding, these offsprings became profoundly hyperinsulinemic, insulin-resistant and glucose-intolerant compared to controls from euthyroid mothers. Suggesting presence of epigenetic changes, altered glucose metabolism was maintained in a second generation of animals. As such, gestational hypothyroidism induces long-term and persistent alterations in endocrine pancreas function, which may have important implications for type 2 diabetes prevention in affected individuals.

**SIGNIFICANCE:** Diabetes and hypothyroidism are two major public health issues, affecting ∼ 9 and 2 % of the population worldwide, respectively. As master metabolic gatekeepers, the thyroid hormones play an essential role in metabolism and fetal development. However, gestation increases demand on the thyroid axis in the mother, leading to hypothyroidism in 0.5 % of pregnancies. Maternal hypothyroidism is associated with deficits in fetal growth that may lead to long-term alterations in metabolism in the offspring. We therefore sought to investigate the effects of gestational hypothyroidism on glucose metabolism in adult offspring and their descendants, and how this may predispose to diabetes development.

## INTRODUCTION

Type 2 diabetes (T2D) and hypothyroidism are two major public health issues, affecting ∼ 9 % and 2 %, respectively, of the population worldwide (1, 2). These endocrine pathologies alter whole body metabolism and can be sometimes related, without presenting a common etiology (3, 4). T2D arises from a complex interplay between genetic and environmental factors (5). In particular, the fetal environment plays a key role in the establishment of a functional beta cell mass (6). Changes in the intra-uterine milieu can modify beta cell differentiation and proliferation in the fetus, leading to long-term effects on glucose metabolism (7).

Different maternal conditions alter the circulating levels of nutrients and hormones, which might impact beta cell development in utero. First, pre-existing metabolic disorders, such as malnutrition, obesity and diabetes, have been linked to increased susceptibility of the offsprings to chronic diseases, such as hypertension and diabetes (7, 8). In mice, maternal diabetes induces fetal hyperglycemia and hyperinsulinemia through accelerated endocrine pancreas development, predisposing to T2D at later stages (9). Second, gestation itself leads to important metabolic and hormonal modifications. For instance, gestational diabetes, occurring in 13 % of pregnancies (10), alters endocrine pancreas maturation in the fetus and constitutes a risk factor for T2D in adulthood (9). In addition, gestation increases demand on thyroid hormones in the mother, leading to hypothyroidism in 0.5 % of pregnancies (11).

As master metabolic gatekeepers, the thyroid hormones thyroxine (T4) and triiodo-thyronine (T3) play an essential role in metabolism and fetal development. Maternal hypothyroidism is associated with deficits in fetal growth and cardiac, nervous and bone maturation (12, 13). Such dramatic effects result from a complete dependence of the fetus on maternal thyroid hormones until mid-gestation in mice and second trimester of pregnancy in humans (14, 15), and a continued influence of maternal thyroid hormones at later stages (14). At the level of the pancreas, different studies have demonstrated important effects of thyroid hormones on beta cell development and maturation (16, 17). These effects can be direct, through specific interactions with cognate receptors on beta cells (18), or indirect, through modification of the availability of growth factors (16, 19), thereby altering glucose metabolism and insulin resistance (3). Although a recent study showed that fetal hypothyroidism in sheep leads to increased beta cell proliferation and hyperinsulinemia in the fetus (17), the consequences of gestational hypothyroidism on beta cell function in adult offsprings remains unexplored. Thus, we sought to investigate the effects of gestational hypothyroidism on beta cell maturity and function, glucose metabolism, and susceptibility to metabolic stress such as high-fat diet (HFD) in adult mice offsprings and their descendants.

## RESULTS

### Maternal hypothyroidism alters glucose homeostasis in adult offsprings

Gestational hypothyroidism in female mice was induced from the first day post-coitus by administration of an iodine-deficient diet supplemented with propylthiouracil (PTU), known to block thyroid hormone synthesis and conversion (20). This approach led to severe hypothyroidism, shown by a marked post-partum decrease in total T4 hormone levels (Fig. 1A), as previously described (20, 21). Although male mice born to hypothyroid mothers did not display significant changes in weight gain compared to control mice (Fig. 1B) and were euthyroid (Fig. 1C), glucose metabolism in adults (8-10 weeks) was altered. In particular, male offsprings of iodine-deficient mothers presented higher fasting blood glucose levels (Fig. 1D) but improved glucose tolerance (Fig. 1E).Insulin sensitivity was also improved (Fig. 1F), depicted by increased fasting insulin levels (Fig. 1G) and lower glucose-stimulated insulin release during intraperitoneal glucose tolerance test (IPGTT) (Fig. 1H).

**Figure 1.**
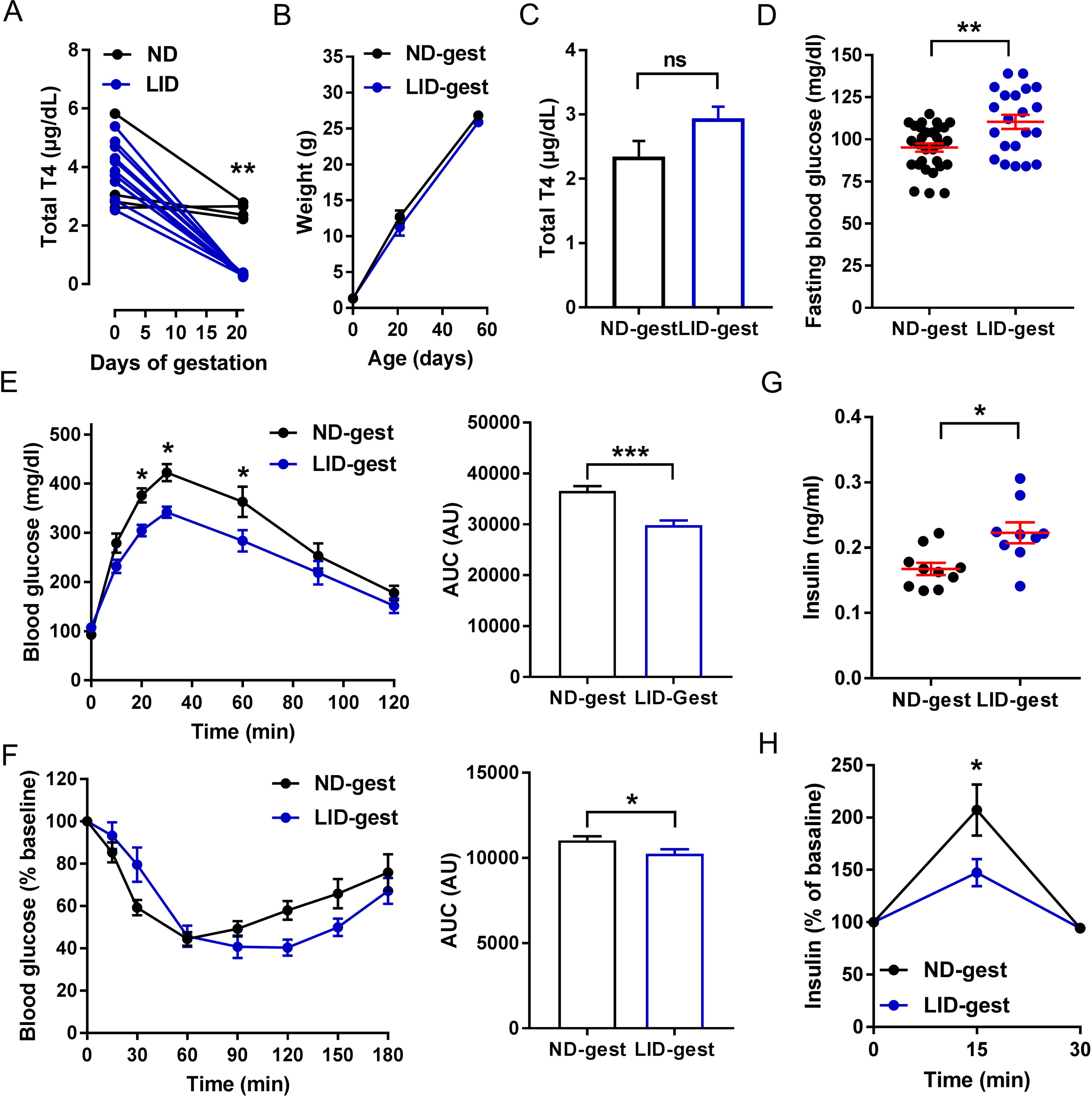
Congenital hypothyroidism alters glucose metabolism in male offspring. ND: normal diet (in black), LID: low-iodine diet (in blue), gest: gestation. Adult male offsprings (8-10 weeks of age) were analyzed. A) Total plasma T4 concentration in dams before gestation and postpartum (n = 4-11 mice/group, mean ± SEM, Mann-Whitney). B) Male offspring weight over time after birth (n = 15-38 mice/group, mean ± SEM, One-way ANOVA). C) Total plasma T4 concentration in adult male offsprings (n = 5-7 mice/group, mean ± SEM, Mann-Whitney). D) Fasting blood glucose in adult male offsprings (n = 21-31 mice/group, mean ± SEM, Mann-Whitney). E) Intra-peritoneal glucose tolerance test (IPGTT) using 3 g glucose/kg body weight (n = 10 mice/group, mean ± SEM, Two-way ANOVA) and area under the curve (AUC) analysis) (n = 10 mice/group, mean ± SEM, Mann-Whitney). F) Insulin tolerance test (ITT (0.75 U insulin/kg) and AUC analysis) (n = 8-10 mice/group, mean ± SEM, Mann-Whitney). G) Fasting basal insulin levels (n = 10 mice/group, mean ± SEM, Mann-Whitney). H) Insulin secretion during IPGTT (3 g/kg), (n = 9 mice/group, mean ± SEM, One-way ANOVA).

In female offsprings, alterations in glucose metabolism were also observed, albeit less pronounced (Fig. 2). Female mice born to hypothyroid mothers presented similar weight gain from birth to adulthood compared to control mice (Fig. 2A), and were euthyroid (Fig. 2B). Although fasting blood glucose levels were similar to those in control mice (Fig. 2C), female offsprings of iodine-deficient mothers displayed improved glucose tolerance (Fig. 2D). However, insulin sensitivity (Fig. 2E), fasting insulin levels (Fig. 2F) and glucose-stimulated insulin release (Fig. 2G) remained unchanged.

**Figure 2.**
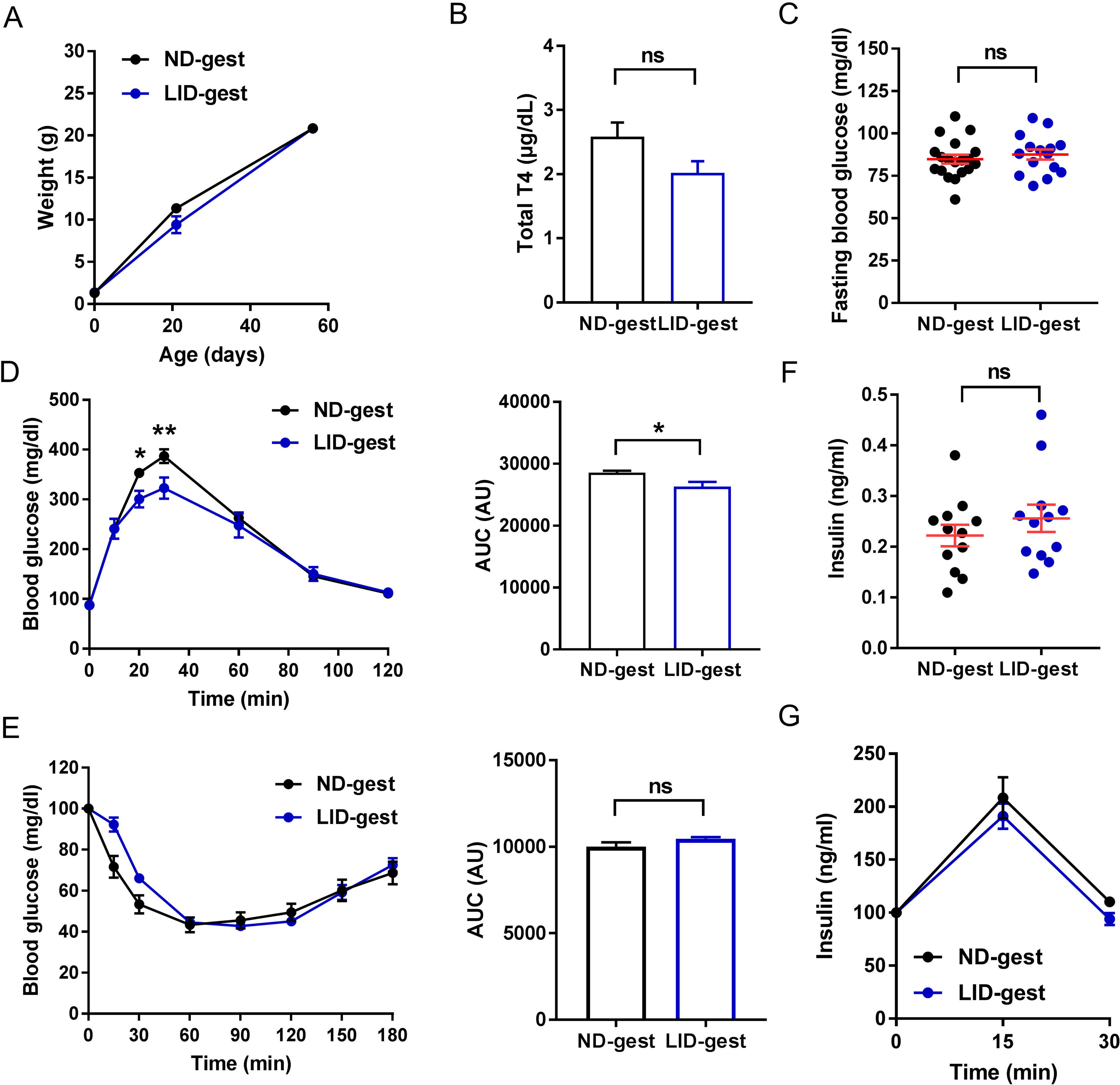
Congenital hypothyroidism alters glucose metabolism in female offspring. ND: normal diet (in black), LID: low-iodine diet (in blue), gest: gestation. Adult female offspring (8-10 weeks of age) were analyzed. A) Female offspring weight over time after birth (n = 15-19 mice/group, mean ± SEM, One-way ANOVA). B) Total plasma T4 concentration in adult female offspring (n = 5-10 mice/group, mean ± SEM, Mann-Whitney). C) Fasting blood glucose in adult female offspring (n = 21-31 mice/group, mean ± SEM, Mann-Whitney). D) Intra-peritoneal glucose tolerance test (IPGTT (3 g/kg) (n = 10 mice/group, mean ± SEM, Two-way ANOVA) and area under the curve (AUC) analysis) (n = 10 mice/group, mean ± SEM, Mann-Whitney). E) Insulin tolerance test (ITT (0.75 UI/kg) and AUC analysis) (n = 13-14 mice/group, mean ± SEM, Mann-Whitney). F) Fasting insulin levels (n = 12 mice/group, mean ± SEM, Mann-Whitney). H) *In* vivo insulin responses to glucose (3 g/kg), (n = 11 mice/group, mean ± SEM, One-way ANOVA).

### Gestational hypothyroidism alters beta cell proliferation

Morphometric analyses of pancreatic sections showed that islet area was similar in offsprings born to hypothyroid and euthyroid mothers (Fig. 3A-B). However, Ki67 labeling in beta cells, a marker of proliferation, was found to be ∼2-fold higher both in males and females born to hypothyroid mothers (Fig. 3A, C).

**Figure 3.**
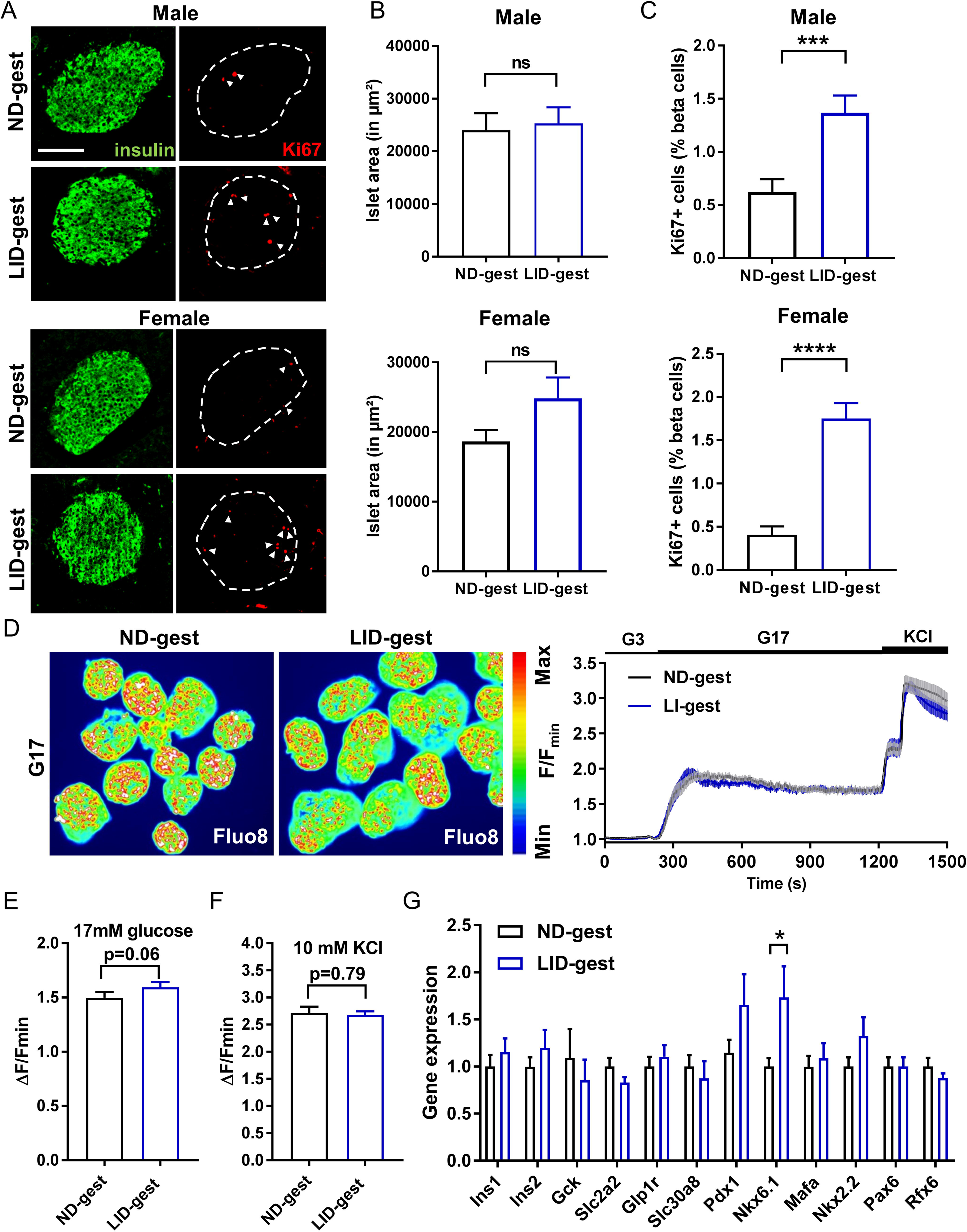
Gestational hypothyroidism alters beta cell proliferation without changes in and calcium fluxes and beta cell identity. ND: normal diet (in black), LID: low-iodine diet (in blue), gest: gestation. Adult offspring (8-10 weeks of age) were analyzed. A) Confocal images of pancreatic islets in male (top panels) and female (bottom panels) offspring (scale: 100 µm, 5 µm Z-projection; red: Ki67, green: insulin). Dashes circles delineate islets and arrows indicate Ki67+ beta cell nuclei. B) Quantification of islet area in male (top panel) and female (bottom panel) offspring (n = 5-12 mice/group, mean ± SEM, Mann-Whitney). C) Quantification of beta cell proliferation in male (top panel) and female (bottom panel) offsprings (measured as % of beta cells positive for Ki67) (n = 5-12 mice/group, mean ± SEM, Mann-Whitney). D-F) Representative images (D, left), mean traces (D, right) and summary bar graphs (E and F) showing no changes in the amplitude of glucose and glucose + KCl-stimulated Ca^2+^ rises in LI-gest male offspring (n = 44-61 islets/9-12 mice, mean± SEM, Mann-Whitney). G) Expression profiles of key beta cell markers in islets from male offsprings, measured by RT-qPCR. Data were normalized by the geometric mean of *Ppia* and *Mrlp32.* Ct values are expressed as fold increase relative to offspring from ND-fed control. Data is presented as mean ± SEM (n = 10 mice/group, Mann-Whitney).

Suggesting that defects in insulin secretion are distal to the triggering pathway, glucose- and KCl-stimulated Ca^2+^ rises were unchanged in islets from male animals born to hypothyroid mothers (Fig. 3D-F). Furthermore, the islet mRNA abundance of genes responsible for maintenance of beta cell identity and maturity remained unchanged (*Mafa*, *Pdx1*, *Rfx6*, *Pax6, Ins1*, *Ins2, Slc2a2, Gck and Glp1r*), with the exception of increased *Nkx6-*1 mRNA levels (Fig. 3G).

### Gestational hypothyroidism renders offsprings more susceptible to metabolic stress

We next explored whether gestational hypothyroidism would influence compensatory responses to metabolic stress on adult male offsprings. Animals born to hypothyroid mothers displayed increased weight gain following high fat diet feeding (Fig. 4A-B), whereas weight gain was comparable when animals were on normal chow for the same duration (Fig. 4A-B). While HFD increased fasting glucose (Fig. 4C), and induced glucose intolerance (Fig. 4D) in all animals examined, the defect was most severe in offsprings born to hypothyroid mothers, despite similar insulin sensitivity to age-matched controls (Fig. 4E). A similar trend was seen in fasting insulin levels (Fig. 4F), with pronounced hyperinsulinemia present in offspring born to hypothyroid mothers, despite the absence of change in insulin levels following glucose challenge (Fig. 4G).

**Figure 4.**
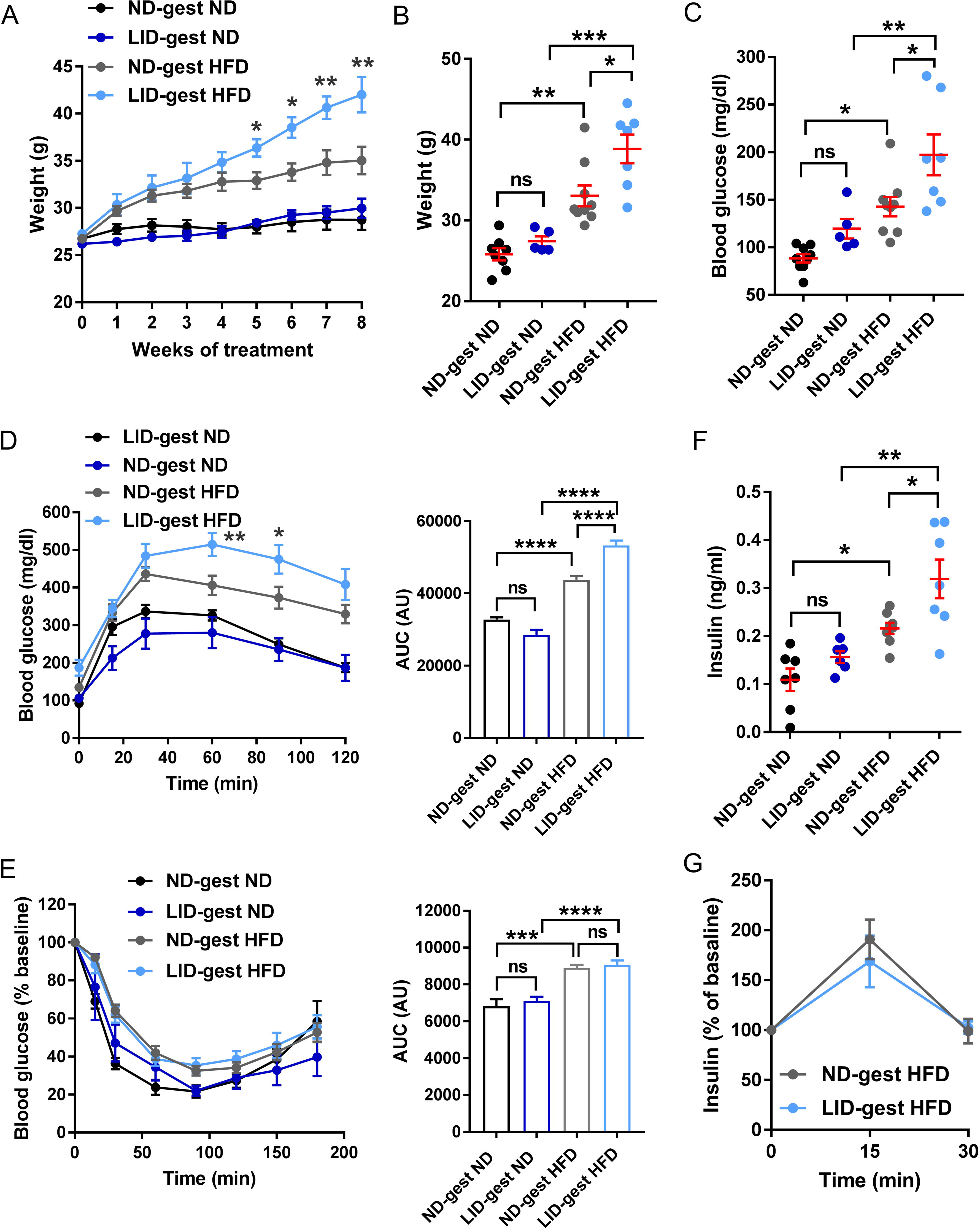
Gestational hypothyroidism renders offspring more susceptible to metabolic stress. ND-gest ND: normal diet during gestation then normal diet (in black), LID-gest ND: low-iodine diet during gestation then normal diet (in dark blue), ND-gest HFD: normal diet during gestation then high fat diet (in grey), LID-gest HFD: low-iodine diet during gestation then high fat diet (in light blue). Adult male offspring (8-10 weeks of age) were analyzed. A) Growth curve post initiation of feeding treatments showing increased weight gain in male offsprings from hypothyroid mothers (n = 5-10 mice/group mean ± SEM, Two-way ANOVA). B) Animal weight at the end of feeding treatment (8 weeks) (n = 5-10 mice/group, mean ± SEM, One-way ANOVA). C) Fasting blood glucose at the end of feeding treatment (8 weeks) (n = 5-10 mice/group, mean ± SEM, One-way ANOVA). D) Intra-peritoneal glucose tolerance test (IPGTT (3 g/kg) (n = 5-8 mice/group, mean ± SEM, Two-way ANOVA) and area under the curve (AUC) analysis) (n = 5-8 mice/group, mean ± SEM, One-way ANOVA). E) Insulin tolerance test (ITT (0.75 UI/kg) and AUC analysis) (n = 5-8 mice/group, mean ± SEM, One-way ANOVA). F) Fasting basal insulin levels (n = 5-10 mice/group, mean ± SEM, One-way ANOVA). G) *In* vivo insulin responses to glucose (3 g/kg), (n = 8 mice/group, mean ± SEM, One-way ANOVA).

At the morphological level, HFD increased islet size and beta cell proliferation to a similar extent in offsprings from both hypothyroid or euthyroid mothers (Fig. 5A-C), as expected (22). Using multicellular live Ca^2+^ imaging of intact islets (23), no differences in glucose- or KCl-stimulated Ca^2+^ rises were detected between the two groups of animals (Fig. 5D-F), suggesting that HFD does not affect preferentially ionic fluxes or glucose-responsiveness in males born to hypothyroid mothers.

**Figure 5.**
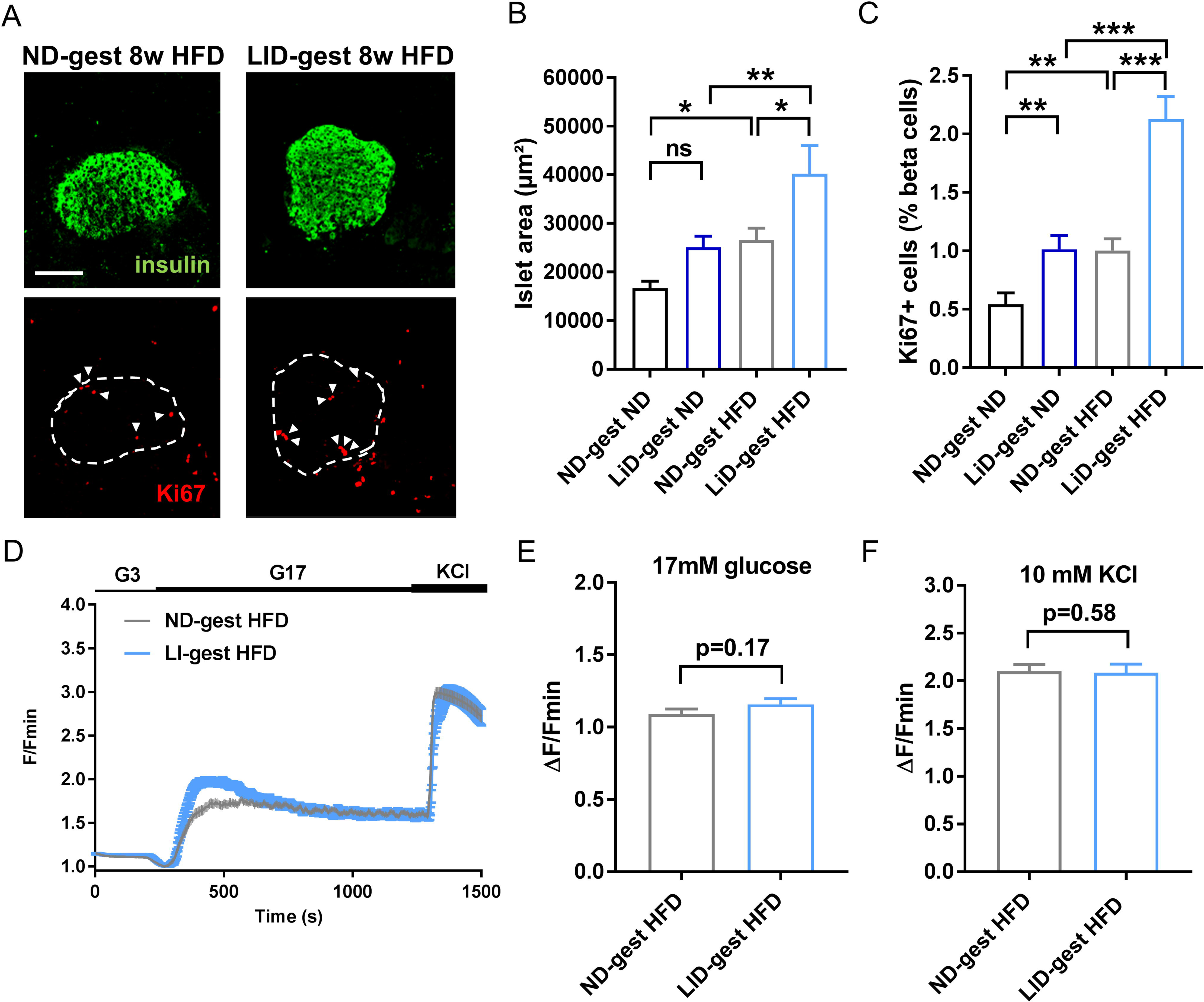
Metabolic stress-induced changes in beta cell proliferation and calcium fluxes are comparable between offspring from hypothyroid and euthyroid mothers. ND-gest ND: normal diet during gestation then normal diet (in black), LID-gest ND: low-iodine diet during gestation then normal diet (in dark blue), ND-gest HFD: normal diet during gestation then high fat diet (in grey), LID-gest HFD: low-iodine diet during gestation then high fat diet (in light blue). Adult male offspring (8-10 weeks of age) were analyzed. A) Confocal images of pancreatic islets in male offspring (scale: 100 µm, 5 µm Z-projection; red: Ki67, green: insulin). Dash circles delineate islets and arrows indicate Ki67+ beta cell nuclei. B) Quantification of islet area (n = 5-12 mice/group, mean ± SEM, One-way ANOVA). C) Quantification of beta cell proliferation (measured as % of beta cells positive for Ki67) (n = 5-12 mice/group, mean ± SEM, One-way ANOVA). D-F) Mean traces (D) and summary bar graphs (E and F) showing no changes in the amplitude of 17 mM glucose and glucose + 10 mM KCl-stimulated Ca^2+^ rises in LID-gest offspring fed HFD (n = 58-76 islets/6-7 mice/group, mean ± SEM, Mann-Whitney).

### The effects of gestational hypothyroidism on glucose homeostasis are transgenerational

We finally investigated whether alterations in glucose metabolism persisted in a second generation of animals. To do this, glucose tolerance, insulin resistance, fasting insulin levels and beta cell proliferation were assessed in adult offspring (8-10 weeks of age) originating from male mice born to hypothyroid mothers. While the second generation of males presented similar weight and fasting blood glucose compared to control aged-matched animals (Fig. 6A-B), glucose tolerance remained improved (Fig. 6C). This change was despite normal insulin sensitivity (Fig. 6D), decreased fasting insulin levels (Fig. 6E) and normal glucose-stimulated insulin release (Fig. 6F). Second generation female offsprings also presented similar weight compared to control aged-matched animals (Fig. 6G). By contrast to males, glucose tolerance was unchanged (Fig. 6I), despite increased fasting blood glucose (Fig. 6H) and decreased insulin sensitivity (Fig. 6J), decreased fasting insulin levels (Fig. 6K) and normal glucose-stimulated insulin release Fig. 6L). Finally, islet size and beta cell proliferation were unchanged in both male and female offspring (Fig. 6M-P). Thus, gestational hypothyroidism induces transgenerational changes in metabolism, with differential effects in male and female offspring.

**Figure 6.**
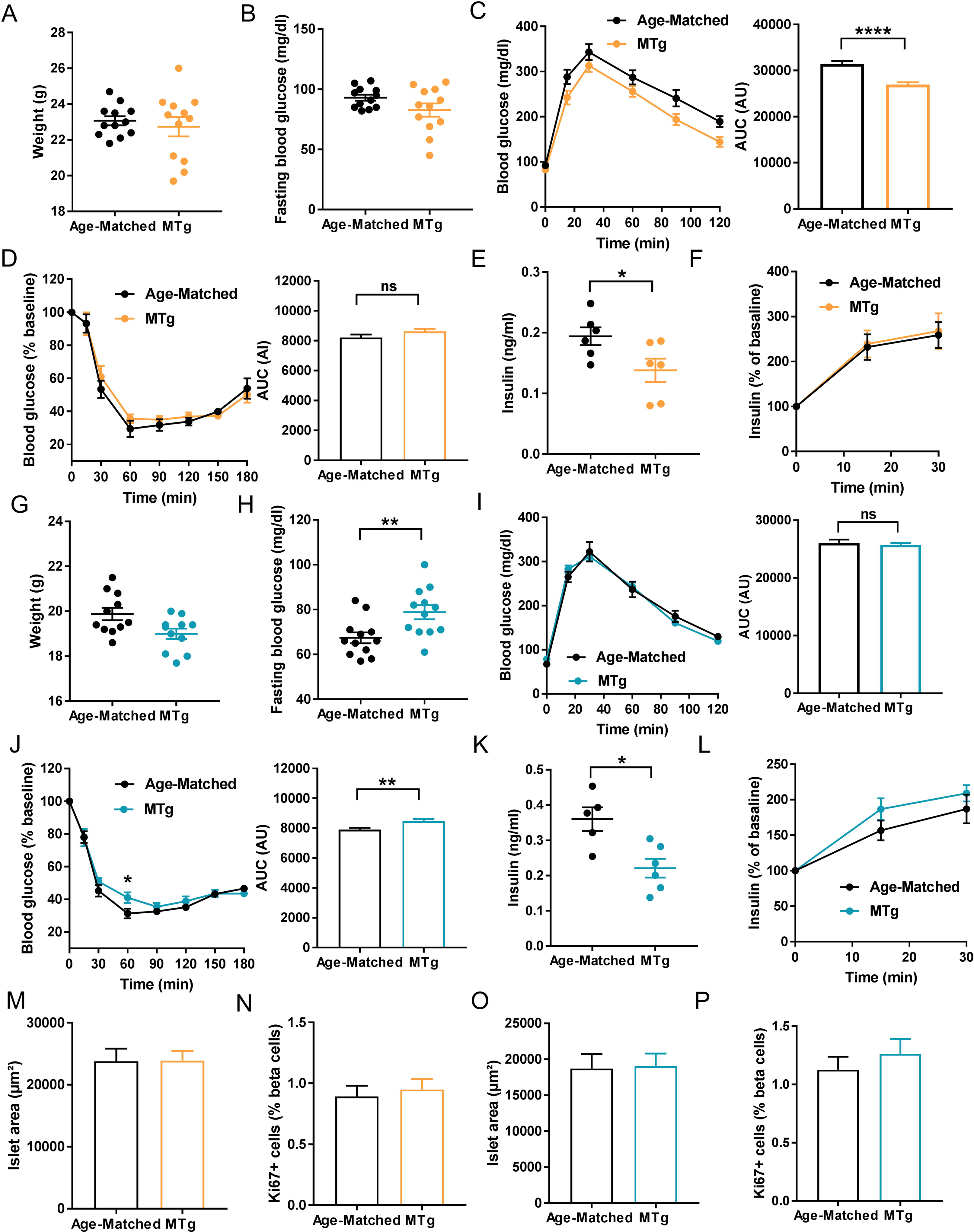
Effects of gestational hypothyroidism on glucose homeostasis in a second generation of animals. MTg: second generation of male (in orange) and female (in cyan) offspring from males born to hypothyroid mothers. Adult MTg (8-10 weeks of age) were compared to age-matched controls (in black). A) Male weight at adult age (8-10 weeks) (n = 12 mice/group, mean ± SEM, Mann-Whitney). B) Fasting blood glucose in males (n = 12 mice/group, mean ± SEM, Mann-Whitney). C) Glucose tolerance test in males (3 g/kg) and area under the curve (AUC) analysis (n = 12 mice/group, mean ± SEM, Mann-Whitney). D) Insulin tolerance test in males (0.75 UI/kg) and AUC analysis (n = 7-10 mice/group, mean ± SEM, Mann-Whitney). E) Fasting insulin levels in males (n = 6 mice/group, mean ± SEM, Mann-Whitney). F) *In* vivo insulin responses to glucose in males (3 g/kg), (n = 6 mice/group, mean ± SEM). G) Female weight at adult age (8-10 weeks) (n = 12 mice/group, mean ± SEM, Mann-Whitney). H) Fasting blood glucose in females (n = 12 mice/group, mean ± SEM, Mann-Whitney). I) Glucose tolerance test in females (3 g/kg) and area under the curve (AUC) analysis (n = 12 mice/group, mean ± SEM, Mann-Whitney). J) Insulin tolerance test in females (0.75 UI/kg) and AUC analysis (n = 7-10 mice/group, mean ± SEM, Two-way ANOVA (left panel) and Mann-Whitney (right panel)). K) Fasting insulin levels in females (n = 5-6 mice/group, mean ± SEM, Mann-Whitney). L) *In* vivo insulin responses to glucose in females (3 g/kg), (n = 5-6 mice/group, mean ± SEM). M) Quantification of islet area in males (n = 6 mice/group, mean ± SEM, Mann-Whitney). N) Quantification of beta cell proliferation in males (n = 6 mice/group, mean ± SEM, Mann-Whitney). O) Quantification of islet area in females (n = 6 mice/group, mean ± SEM, Mann-Whitney). P) Quantification of beta cell proliferation in females (n = 6 mice/group, mean ± SEM, Mann-Whitney).

## DISCUSSION

Circulating factors *in utero* can influence fetal endocrine pancreas development and lead to life-long alterations in glucose metabolism. Since gestation modulates thyroid hormone levels which are known to play an important role in beta cell development and maturation (12, 17), we sought to investigate whether maternal hypothyroidism influences glucose homeostasis in adult offspring. We found that gestational hypothyroidism induced by iodine-deficient diet increased beta cell proliferation, altered glucose metabolism and increased severity of high-fat diet-induced obesity in the offspring, without altering beta cell maturity and functional responses. Furthermore, alterations in glucose metabolism were maintained in a second generation of adults. We thus provide evidence that maternal hypothyroidism exerts transgenerational effects on metabolism, manifested by glucose intolerance in response to metabolic stress.

Hypothyroidism is one of the most common endocrine diseases during pregnancy and is mainly linked to dietary iodine deficiency, especially in low-middle income countries (3). Thus, iodine deficiency in diet constitutes a reliable model to induce congenital hypothyroidism through severe decreases in circulating total T4 levels during gestation (24). However, corresponding studies in mice remain scarce. While exposure to hypothyroidism *in utero* has been reported to influence growth in other rodents (25, 26), we could not detect significant differences in body mass between mice born to euthyroid or hypothyroid mothers. While this may reflect the model used, we note that intrauterine growth restriction does not necessarily correlate with altered body weights in neonates (27). Indeed, in sheep, hypothyroidism in utero induced pancreatic beta cell proliferation and hyperinsulinaemia in the fetus (17), which would be expected to maintain growth rate.

Although effects of thyroid hormone deficiency on fetal pancreas development were not assessed here, congenital hypothyroidism in mice altered glucose metabolism and stimulated beta cell proliferation in both adult male and female mouse offsprings. This suggests impaired development of the fetal or postnatal pancreas, leading to long-term alterations in glucose metabolism (28). Overall, the alterations were more pronounced in male offspring, possibly reflecting known sex-dependent effects of fetal hypothyroidism on glucose metabolism (29). In line with our results, congenital hypothyroidism has been previously shown to induce long-term alterations in glucose metabolism in adult male rat offsprings (28, 30). In particular, insulin secretion was found to be decreased, in contrast to the increase detected in the present study. The reasons for this are unknown, but may include the dynamic evolution of glucose metabolism with age (young adult versus mature animals), the proliferative status of beta cells (not assessed in rat studies), and/or species-related differences.

Since glucose-stimulated Ca^2+^ fluxes are a major triggering signal for insulin release (31), we hypothesized that long-term alterations in glucose metabolism induced by congenital hypothyroidism might be linked to changes in beta cell stimulus-secretion coupling. However, both glucose- and KCl-induced Ca^2+^ rises were found to be unchanged in islets from male animals born to hypothyroid mothers. Previous studies on isolated islets from male rat offspring showed that maternal hypothyroidism led to impaired insulin secretion through a combination of different mechanisms, including alteration in glycolytic pathways and K_ATP_ and L-type Ca^2+^ channel conductance (30). Thus, the changes in *in* vivo insulin responses and glucose metabolism described in the present study likely result from mechanisms distal to Ca^2+^ fluxes, such as amplyfying pathway (e.g. cAMP) or granule exocytosis. Alternatively, since thyroid hormones are crucial regulators of growth, development and metabolism in virtually all tissues, in particular during fetal stages (12), metabolic alterations may rise from a combination of modifications in different organs. For instance, congenital hypothyroidism has been shown to alter liver development (32), and modify glucose transporter expression, impairing glucose sensing in glucose-sensitive organs, including the liver and metabolic brain (33). Further studies will be needed to explore both these possibilities.

In adults, beta cell proliferation is triggered in response to increased metabolic demand such as gestation and high-fat diet feeding (22, 34). Although triiodothyronine stimulates proliferation of rat beta cell lines (35), whether thyroid hormones contribute to beta cell proliferation in response to demand remains unclear. During fetal development, the prepartum surge in thyroid hormone is thought to induce a switch from beta cell proliferation to functional maturation (16, 36), thus explaining the maintenance of beta cell proliferation in islets of hypothyroid sheep fetuses (17). While similar mechanisms are likely at play in offspring of hypothyroid mothers, we cannot exclude an increase in beta cell proliferation due to increased metabolic demand. The source of such increased demand is however unclear, especially since normal diet-fed offspring displayed increase insulin sensitivity. Since T3/T4 are pre-requisite for cell maturation (37), and because *in vivo* insulin response to glucose was decreased, we analyzed overall gene expression of key markers defining adult beta cell functional identity (38). However, we could not detect major changes in adult offspring from hypothyroid mothers, suggesting that beta cell de-differentiation/or lack of maturation is not a feature here.

In addition to altered glucose metabolism and increased beta cell proliferation, maternal hypothyroidism increased susceptibility to HFD-induced metabolic stress in adult male offspring. This is in line with previous data showing that alterations in endocrine pancreas development can induce long-term consequences for glucose metabolism (7, 39). The results here support that maternal hypothyroidism may increase risk of type 2 diabetes development in later life. This increased susceptibility may be linked to exacerbated HFD-induced hyperinsulinemia, which has previously been shown to drive insulin resistance and diet-induced obesity (40). Again, changes were independent of Ca^2+^ channel activity, suggesting that insulin secretory defect may lie distal to the triggering pathway. Although female offspring were not analyzed, similar results may likely be obtained, since congenital hypothyroidism also affected glucose metabolism in female offspring.

Finally, we saw that altered glucose metabolism persisted in a second generation of offspring, albeit to a lesser extent, suggesting the presence of epigenetic changes. Such changes are likely to be imprinted due to thyroid hormone deprivation during fetal development, since epigenetic reprogramming occurs during gametogenesis and early embryogenesis (41), before being transmitted to the next generation. Since both liver and pancreas are affected by similar signaling pathways during development and both organs display remarkable plasticity following insult in adults, epigenetic markers likely affect other organs than the endocrine pancreas (42). We concede, however, that identification of these epigenetic markers is needed, and that a multitude of other mechanisms may also be involved in altered glucose homeostasis following changes in thyroid hormone levels.

In summary, we show that gestational hypothyroidism induces trans-generational effects on glucose metabolism in the offspring, which may affect predisposition to T2D development in response to metabolic stress.

## MATERIAL AND METHODS

### Mice

Animal studies were conducted according to the European guidelines for animal welfare (2010/63/EU). Protocols were approved by the Institutional Animal Care and Use Committee (CEEA-LR-1434) and the French Ministry of Agriculture (APAFIS#13044). Mice were housed in conventional facility on a 12 h light-12 h dark cycle and were given chow and water *ad libitum*. FVB mice were purchased from Janvier-SAS (Le-Genest-St-Isle, France). Hypothyroidism in gestating mice was induced by feeding animals with iodine-deficient diet supplemented with 0.15% propylthiouracil (PTU) (Envigo) (20) from the first day post-coitus. Offspring were subsequently fed with normal diet (ND) until age 8-10 weeks and then fed with either normal diet (ND) or high fat diet (HFD, 63% calories from fat) (Safe Diets) for 8 weeks. Analyzed offspring were from at least three independent breeding pairs per group. Second generation animals originated from two different male breeders born to hypothyroid mothers. Intra-peritoneal glucose tolerance tests (IPGTT), insulin tolerance tests (ITT) and glucose-stimulated insulin secretion *in vivo* (GSIS) tests were as described (43, 44). Mice were euthanized by decapitation after asphyxiation with CO_2_. Total trunk blood was collected and total T4 was measured in plasma in duplicate using a total T4 Elisa kit (EIA-1781, DRG International).

### Confocal imaging and image analysis

Pancreas preparation and antibody labeling were as described (44). Briefly, pancreas were fixed overnight in 4% paraformaldehyde and sliced on a vibratome (Leica) before immune-staining. Antibodies used were: rabbit anti-Ki67 (1:200, CliniSciences), guinea-pig anti-insulin (1:400, Abcam), mouse anti-glucagon (1:200, Sigma). Nuclei were labeled using dapi (Sigma). Images were acquired using a Zeiss LSM 780 confocal microscope. Images were analyzed using Imaris (Bitplane), Volocity (Perkin Elmer) and ImageJ (NIH). For quantifications, four slices were randomly selected from at least three animals/group, and all islets present analyzed. *A priori*, this is sufficiently-powered to detect a minimum 1.2-fold difference with a SD of 40%, a power of 0.9, and alpha = 0.05 (G*Power 3.1). The proportion of proliferative beta cells was obtained by dividing number of Ki67+ nuclei by total number of nuclei of insulin+ cells in islets, as described (44).

### Islet isolation and live calcium imaging of isolated islets

Islets were hand-picked after collagenase digestion of the whole pancreas (45), and cultured (5% CO_2_, 37°C) in RPMI medium containing 10% FCS, 100 units/mL penicillin, and 100 μg/mL streptomycin. Islets were loaded with Fluo-8 (AAT Bioquest) dissolved in DMSO containing 20% pluronic acid. Islets were then imaged using a Crest X-Light spinning disk system coupled to a Nikon Ti-E base and 10 x / 0.4 / air objective. Excitation was delivered at λ = 458–482 nm using a Lumencor Spectra X light engine, with emitted signals detected at λ = 500-550 nm using a Photometrics Delta Evolve EM-CCD. Imaging buffer contained (in mmol/L) 120 NaCl, 4.8 KCl, 24 NaHCO_3_, 0.5 Na_2_HPO_4_, 5 HEPES, 2.5 CaCl_2_, 1.2 MgCl_2_, and 3–17 D-glucose. Data was analyzed using ImageJ, with traces presented as Δ F = fluorescence at any timepoint and F_min_ = minimum fluorescence.

### Islet isolation and real-time quantitative RT-PCR

Pancreatic islets were hand-picked after collagenase digestion of whole pancreas, as described (45). Total RNA from mouse islets was extracted using RNeasy microkit (Qiagen) following the manufacturer’s instructions. Reverse transcription was carried out using random hexamer oligonucleotides and SuperScriptIII Reverse Transcriptase (2,000U; Invitrogen, LifeTechnologies, EUA). The reverse transcription product was diluted according to the efficiency curve and submitted in duplicates to real-time quantitative PCR using LightCycler® 480 SYBR Green I Master (Roche) in 7500 System (Applied Biosystems). Selection of housekeeping genes was performed using NormFinder software (46). PCR reactions were performed following the conditions: 95 °C - 5 min, followed by 45 cycles of 95 °C - 10 s and 72 °C - 30 s and the Melting Curve was performed from 65 up to 97 °C for 1 min. Ct values were recorded for each gene (listed in the Table S1) and normalized to the geometric mean of *Ppia* and *Mrlp32.* Ct values were then expressed relative to offspring from normal diet-fed animals (ND). A list of primers used is shown in Table S1.

### Statistical analysis

Values are represented as mean ± SEM. Statistical tests were performed using GraphPad Prism. Normality was tested using D’Agostino-Pearson test, and comparisons were made using either unpaired Student’s t-test, or two-tailed Mann-Whitney U-test, as appropriate. Multiple comparisons were made using one-way or two-way ANOVA followed by Bonferroni’s post-hoc test. P values were considered significant at P<0.05*, 0.01**, 0.001***, 0.0001****.

## AUTHOR CONTRIBUTIONS

M.S. and D.J.H. conceived and designed experiments; Y.K., D.N., A.D.B., P.B.-S., A.G., and M.S. performed experiments; Y.K., D.N., A.D.B., P.B.-S., R.A.P.G, P.M., D.J.H. and M.S. analyzed data; D.J.H. and M.S. wrote the manuscript.

## ACKNOWLEDGMENTS

The authors declare no conflict of interest. The authors would like to thank Dr. M. Golan, Agricultural Research Organization, Rishon Letziyon, Israel, L. El-Cheik Hussein and E. Galibert, Institute of Functional Genomics, Montpellier, France, for technical assistance, and the animal facility staff (RAM). Authors were supported by grants from the Agence Nationale de la Recherche (ANR BETA-DYN JCJC13 to M.S., France-BioImaging ANR-10-INBS-04 and ANR Peripulse to P.M.), INSERM, CNRS, University of Montpellier, and Région Languedoc-Roussillon. P.B.S. was supported by a São Paulo Research Foundation (FAPESP 2016/24941-7) Fellowship. D.J.H. was supported by a Diabetes UK R.D. Lawrence (12/0004431) Fellowship, a Wellcome Trust Institutional Support Award, MRC Confidence in Concept, MRC (MR/N00275X/1 and MR/S025618/1) Project and Diabetes UK (17/0005681) Project Grants. This project has received funding from the European Research Council (ERC) under the European Union’s Horizon 2020 research and innovation programme (Starting Grant 715884 to D.J.H.).

