## Supplemental Table 1 for "Maternal hypothyroidism in mice influences glucose metabolism in adult offspring"

**SUPPLEMENTAL DATA**

**TABLE S1. List of primers used for quantitative RT-PCR**

| **Symbol**  GenBank# | **Primers** | **Sequences (5’- 3’)** | **Amplicon length (bp)** |
| --- | --- | --- | --- |
| ***Mrlp32***  NM_029271.2 | Forward  Reverse | AGGTGCTGGGAGCTGCTACA  AAAGCGACTCCAGCTCTGCT | 51 |
| ***Ppia***  NM_008907.1 | Forward  Reverse | CAAACACAAACGGTTCCCAG  TTCACCTTCCCAAAGACCAC | 85 |
| ***Ins1***  [NM_008386.4](https://www.ncbi.nlm.nih.gov/nucleotide/NM_008386.4?report=genbank&log$=nuclalign&blast_rank=14&RID=5V8MX9B4014) | Forward  Reverse | CTTCAGACCTTGGCGTTGGA  TGCTGGTGCAGCACTGATCC | 67 |
| ***Ins2***  NM_001185083.2 | Forward  Reverse | CATCAGCAAGCAGGAAG  CCCAGAGGAAGAGCAG | 83 |
| ***Gck***  NM_010292.5 | Forward  Reverse | CTTCACCTTCTCCTTCCCTG; ATCTCAAAGTCCCCTCTCCT | 150 |
| ***Pax6***  NM_001244198.2 | Forward  Reverse | TGAGAAGTGTGGGAACCAGC; AAGTCTTCTGCCTGTGAGCC | 70 |
| ***Nkx2.2***  NM_001077632 | Forward  Reverse | GCCTCCAATACTCCCTGC; GGTCTCCTTGTCATTGTCCG | 110 |
| ***Slc2a2***  NM_031197.2 | Forward  Reverse | CCCTGTTCCTAACCGGGATG; TCCAGGCGAATTTATCCAGCA | 87 |
| ***Nkx6.6***  NM_144955.2 | Forward  Reverse | CGAACAAACGAAGTACTTGGC; TTTCTCCACTTGGTCCTGC | 114 |
| ***Mafa***  NM_194350.1 | Forward  Reverse | GAAGTGCCAGCTCCAGAG; CGCCAACTTCTCGTATTTCTCC | 100 |
| ***Glp1r***  NM_021332.2 | Forward  Reverse | CCCATGGGGGATTGTCAAGT; AGAAAGTTGACGCCGATAGCA | 120 |
| ***Slc30a8***  NM_172816.4 | Forward  Reverse | CGCCTTTTGTATCCTGATTACC; GTTGTAGCCAAAGTTCCGTTG | 120 |
| ***Pdx1***  NM_008814 | Forward  Reverse | CCCTTTCCCGTGGATGAAATC; GAATTCCTTCTCCAGCTCCAG | 145 |
| ***Rfx6***  NM_001159389.1 | Forward  Reverse | TGCTTACCTGCTGGCTGAAA; TGCATTTCTGATTTAACTCCCACC | 96 |
